# Bridging cell-scale simulations and radiologic images to explain short-time intratumoral oxygen fluctuations

**DOI:** 10.1101/2021.03.18.435990

**Authors:** Jessica L. Kingsley, James R. Costello, Natarajan Raghunand, Katarzyna A. Rejniak

## Abstract

Radiologic images provide a way to monitor tumor development and its response to therapies in a longitudinal and minimally invasive fashion. However, they operate on a macroscopic scale (average value per voxel) and are not able to capture microscopic scale (cell level) phenomena. Nevertheless, to examine the causes of frequent fast fluctuations in tissue oxygenation, the models simulating individual cells’ behavior are needed. Here, we provided a link between the average data value recorded for radiologic image voxels and the cellular and vascular architecture of the tissue that fills these voxels. Using hybrid agent-based modeling, we generated a set of tissue morphologies capable of reproducing tissue oxygenation levels observed in radiologic images. We applied this approach to investigate whether oxygen fluctuations can be explained by changes in vascular oxygen supply or by modulations in cellular oxygen absorption. Our studies showed that intravascular changes in oxygen supply can reproduce the observed fluctuations in tissue oxygenation in all considered regions of interest. However, large magnitude fluctuations cannot be recreated by modifications in cellular absorption of oxygen in biologically feasible manner. Additionally, we developed a procedure to identify plausible tissue morphologies for a given temporal series of average data from radiology images. In future applications this approach can be used to generate a set of tissues representative for radiology images and to simulate tumor response to various anti-cancer treatments on the tissue-scale level.

**Authors Summary:** Low levels of oxygen, called hypoxia, are observable in many solid tumors. They are associated with more aggressive malignant cells which are resistant to chemo-, radio- and immunotherapies. Recently developed imaging techniques provide a way to measure the magnitude of frequent short-term oxygen fluctuation, however they operate on a macro-scale voxel level. To examine the causes of rapid oxygen fluctuations on the cell level, we developed a hybrid agent-based mathematical model. We tested two different mechanisms that could be responsible for these cyclic effects in tissue oxygenation: variations in vascular influx of oxygen and modulations in cellular oxygen absorption. Additionally, we developed a procedure to identify plausible tissue morphologies from data collected from radiological images. This will also provide a bridge between the micro-scale simulations with individual cells and the longitudinal medical images containing average voxel values. In the future applications, this approach can be used to generate a set of tissues representative of radiology images and to simulate tumor response to various anticancer treatments on the cell-scale level.

## 1. Introduction

Tumor tissues harbor regions of different levels of oxygen, including the well-oxygenated areas (normoxia) and zones with reduced oxygen availability (hypoxia). The hypoxic regions can arise as a result of rapid proliferation of tumor cells and tortuous tumor vasculature that together lead to an increased distance between some tumor cells and the nearest blood vessel. This, in turn, results in the emergence of oxygen gradients and a diffusion-limited hypoxia, usually at the distances 120-180 µm from vasculature. Such chronic hypoxia may last for a prolonged periods of time, often for more than 24 hours (1, 2). However, hypoxic regions can also be created due to irregular flow of blood in the aberrant tumor vasculature or shut down of small vessels. These phenomena lead to a perfusion-limited hypoxia that is observable for shorter times, often minutes to hours, and can be reversed when the blood flow is restored (1, 2). Several studies demonstrated the existence of 20-30 minutes-long cycles in red blood cell flux that are responsible for periodic changes in oxygen partial pressure (pO_2_) within the tumor tissue (2-4).

However, it has also been observed in murine experiments that tumors experience very fast and sometimes quite large fluctuations in oxygen levels recorded within the tissue. It has been measured using the electron paramagnetic resonance imaging (EPRI) that intratumor oxygen fluctuations can reach a magnitude as high as 40 mmHg over a period as short as 3 minutes (5, 6). An example of this phenomenon recorded in the squamous cell carcinoma VII (SCCVII) tumor is presented in **Figure 1**. In this experiment, four regions of interest (ROIs) were selected based on an anatomical image of the tumor determined using the T2-weighted magnetic resonance imaging (MRI) shown in **Figure 1A**. Each of these regions was characterized by a different initial level of oxygenation from a very well oxygenated Region #1 to a severely hypoxic Region 4. The color-coded map of partial oxygen pressure (pO_2_) is shown in **Figure 1B**. Subsequently, the average pO_2_ values were measured in each region every 3 minutes for 24 minutes (**Figure 1C**). Regions #1 and #2 showed significant changes in oxygenation during this time (more than 10-fold), while Regions #3 and #4 displayed more uniform levels of pO_2_ during the whole experiment. Interestingly, the authors also recorded the levels of EPR tracer during the same time and observed no detectable changes in the tracer levels within each ROI (data not shown here, compare (5, 6)). This indicates that the observed oxygen fluctuations are not related to changes in tissue permeability.

**Figure 1.**
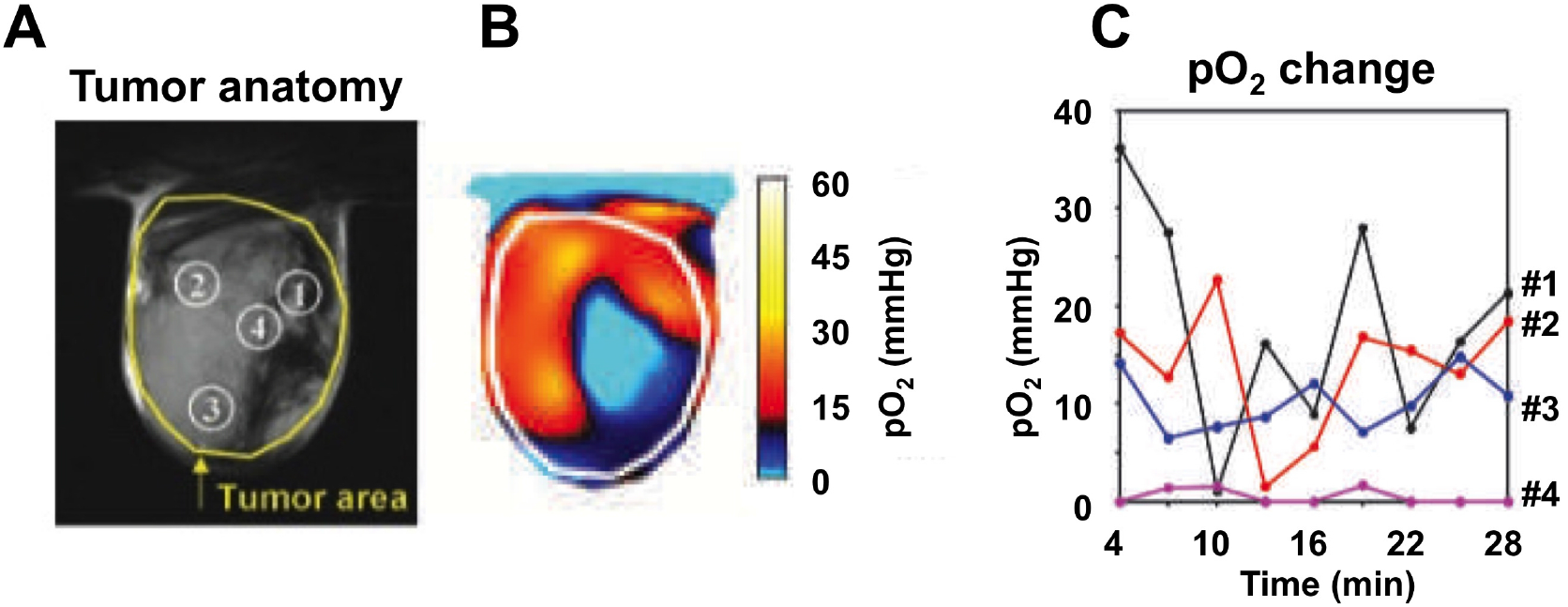
Fast fluctuating pO_2_ in SCCVII squamous cell carcinoma tumors. **A**. T_2_- weighted anatomical image of the tumor with four indicated ROIs when pO_2_ fluctuations were recorded. **B**. A characteristic pO_2_ map obtained by EPRI technique. **C**. The changes in pO_2_ values in each ROI recorded every three minutes (from [6], permission pending).

Our goal is to investigate which mechanisms can lead to these fast and relatively large oxygen fluctuations. In general, the distribution of blood-borne compounds (oxygen, nutrients, drugs, etc.) depends on their vascular supply, interstitial transport and cellular uptake. Since the experimental data indicates that tissue permeability is not a factor in observed fluctuations, we tested whether the changes in vascular oxygen supply or modulations in cellular oxygen absorption can contribute to the observed variations in tissue oxygenation. To address this issue, we used the hybrid agent-based *MultiCell-LF* (multi-cell lattice-free) model (**section 2**) to design a collection of tumor tissues (**section 3.1**) with a stable oxygen gradient (**section 3.2**). For the selected tissues that match the four experimental ROIs (**section 3.3**), we determined optimal rates of oxygen vascular influx or cellular uptake that fit the experimentally observable fluctuations in pO_2_ (**section 3.4**). These optimal schedules were then applied to a larger sample of *in silico* tissues with initial oxygenation levels close to the experimental data to assess schedules’ robustness and reproducibility (**section 3.5**). Finally, we discussed implications of our findings for tumor development and future simulations of tumor treatment (**section 4**).

## 2. Methods – Mathematical Model

For this study, we have considered a tissue patch with an area of 1 mm^2^ that corresponds to a typical voxel in the EPRI oxygen map (7, 8). Tissue morphology and metabolism was modeled using *MultiCell-LF* (9, 10), a hybrid multi-cell lattice-free model that combines the off-lattice individual vessels and cells (tumor and stromal) with a continuous description of oxygen kinetics. A typical example from our simulations is shown in **Figure 2**. The specific pattern of oxygen gradient within the tissue depends on three factors: the amount of oxygen supplied from individual vessels, the amount of oxygen absorbed by both tumor and stromal cells, and the spatial localization of all cells and vessels. As a result of this influx-outflux balance, an irregular gradient of oxygen can emerge.

**Figure 2.**
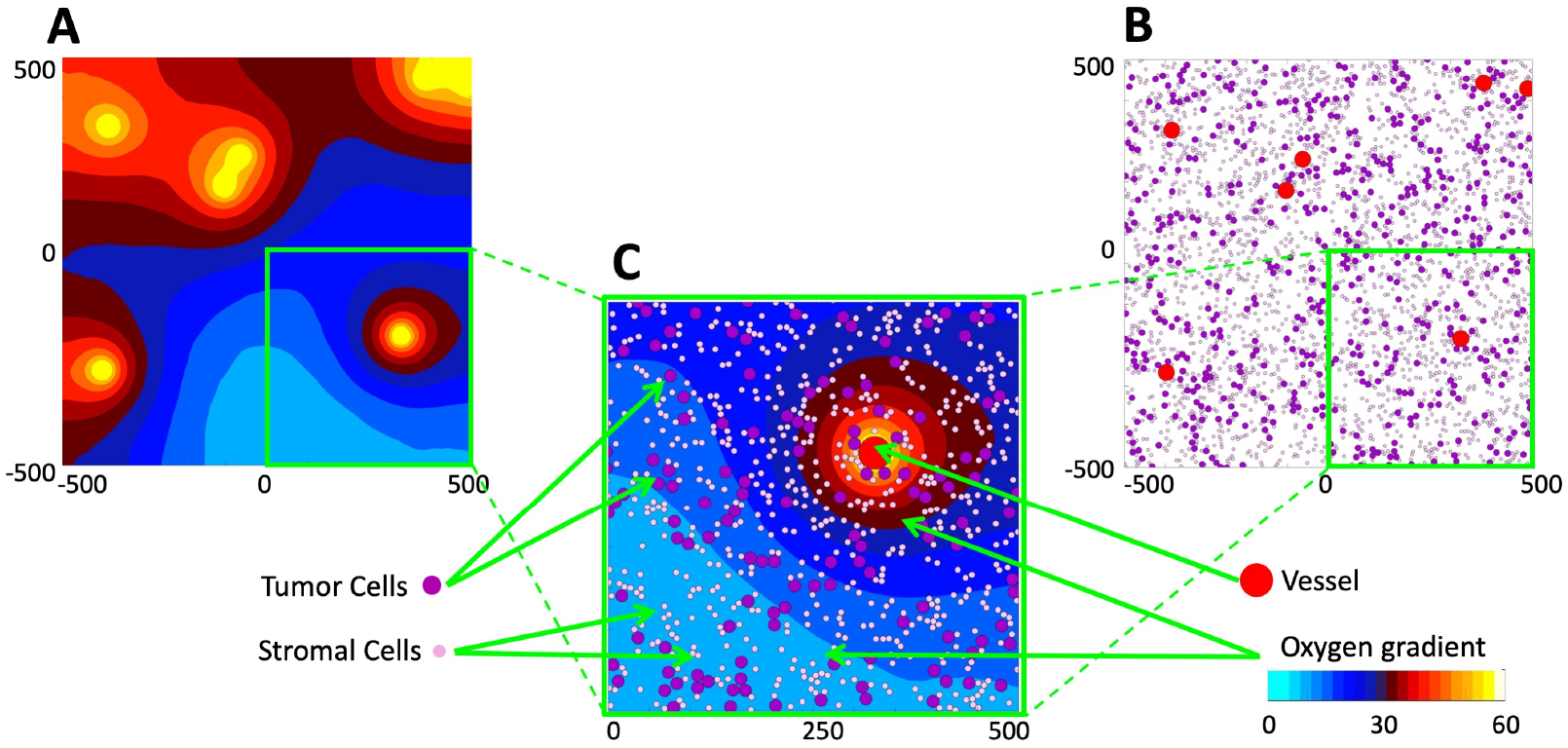
Model of the tumor tissue microenvironment. **A**. A contour map of the gradient of oxygen supplied from the vessels and absorbed by all cells. The color scheme corresponds to that used in EPRI (cyan-low pO_2_ level, white-high pO_2_ level). **B**. Locations of tumor vasculature (red circles), tumor cells (purple circles), and stromal cells (pink circles) within the same computational domain that define tissue cellularity and vascularity. **C**. Magnification of a quarter of the computational domain showing all model components together: the vessels, tumor and stromal cells, and oxygen gradient.

### 2.1 Tissue design

The locations of tumor cells, stromal cells and vessels were chosen randomly within the tissue domain. To ensure that the cells do not overlap with one another and with the vessels, repulsive forces were applied to all cells. Let ***X***_*i*_ and ***X***_*j*_ represent the coordinates of two discrete elements (either tumor cells, stromal cells or vessels) of radii *R*_*i*_ and *R*_*j*_, respectively. The repulsive Hookean force 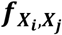 of stiffness *ℱ* acting on element ***X***_*i*_ is given by:

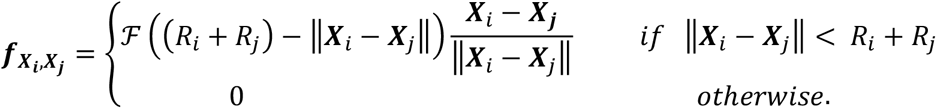

For the tissue that contains *N*_*V*_ vessels of coordinates 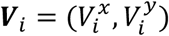, *N*_*T*_ tumor cells of coordinates 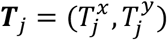, and *N*_*S*_ stromal cells of coordinates 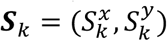, the repulsive forces 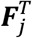 acting on tumor cells and 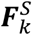 acting on stromal cells combine contributions from all nearby tumor cells, stromal cells and vessels, and are given by the following equations:

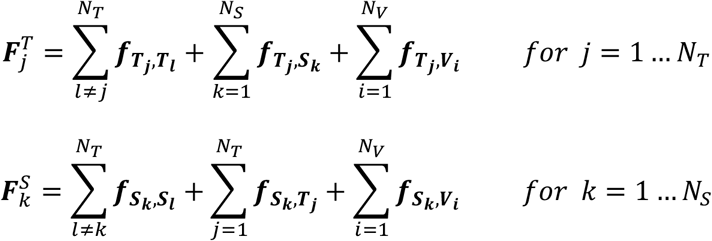

To resolve the overlapping conditions, the tumor and stromal cells are relocated following the overdamped spring equation, where *ν* is the viscosity of the surrounding medium:

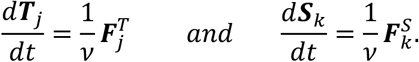

We apply these equations iteratively to all overlapping tumor and stromal cells until the equilibrium is reached with 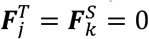. The vessels are not subject to relocation, and we allow the vessels to overlap with other vessels to represent different vascular shapes observed in tissue histologic images. More details on this algorithm are provided in **Supporting Information S1**. Since we will simulate a period shorter than 30 minutes, no cell proliferation and no cell death are included.

### 2.2 Oxygen gradient

Oxygen distribution within the tissue *γ*(***x***, *t*) depends on its influx from vessels, diffusion through the tissue, and the uptake by both stromal and tumor cells. Influx is determined by the location of each individual vessel ***V***_*i*_ and the influx rate *δ*_*V*_(*t*) that can vary over time. The influx rate represents a fraction (0 ≤ *δ*_*V*_(*t*) ≤ 1) of the maximal oxygen supply *γ*_*max*_ characteristic of the oxygen content in the vessels of a given radius *R*_*V*_. The transport through the interstitial space of the tumor tissue is assumed to have a constant diffusion coefficient *𝒟*_*γ*_. The oxygen uptake is defined by the Michaelis-Menten equation with a Michaelis constant *k*_*m*_ common for both tumor and stromal cells, the constant maximum uptake rate for stromal cells *S*_*max*_, and the maximum uptake rate for tumor cells *δ*_*T*_(*t*)*T*_*max*_ for which the rate (*δ*_*T*_(*t*) ≥ 0) can change over time. The kinetics of oxygen is modeled using the following continuous reaction-diffusion equation:

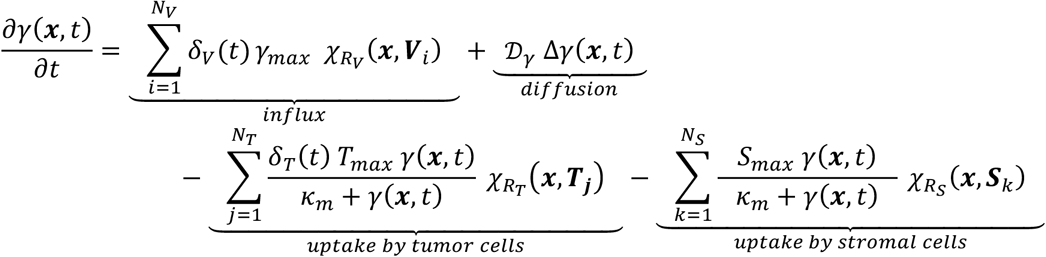

Interactions between oxygen defined on the Cartesian grid ***x*** = (*x, y*) and the tumor cells, stromal cells and vessels defined on the Lagrangian grid ***X*** = (*X, Y*) are specified by the indicator function, *η*_*R*_(***x, X***), with the interaction radius *R*:

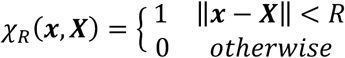

To generate the initial stable oxygen gradient, the oxygen influx rate for each vessel was set up to the maximum vascular level (*δ*_*V*_(*t*) = 1), and tumor cellular uptake was set up to the base uptake level (*δ*_*T*_(*t*) = 1). However, in order to generate oxygen fluctuations in the whole tissue, one or both of these rates were varied. The values of all other model parameters are listed in **Table 1** and more information is provided in **Supporting Information S2**.

**Table 1:**
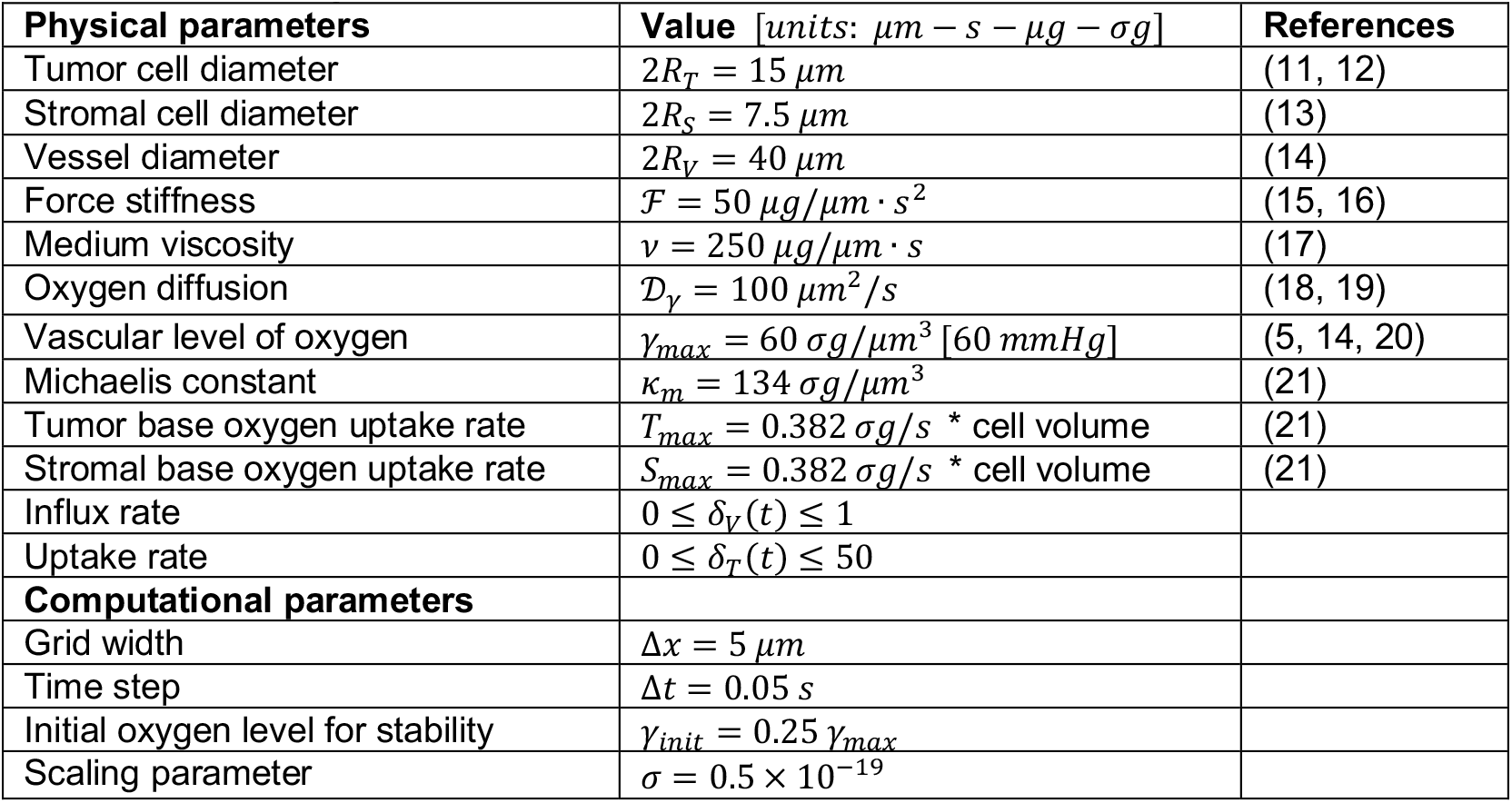
Model physical and computational parameters.

## 3. Results

The goal of this study is to identify the possible mechanisms leading to fast (within 3-minute duration) fluctuations in oxygen level observed in *in vivo* tumors. To do that, we first generated a collection of tumor tissues that varied in the vascular and cellular fractions (**section 3.1**) and determined the stable levels of oxygenation within these tissues (**section 3.2**). Next, we identified *in silico* tissues for which the stabilized oxygenation best fit the four specific experimental data (**section 3.3**). These tissues were used to investigate whether the observed short-time oxygen fluctuations can arise as an effect of altered oxygen supply from the vasculature or altered oxygen uptake by tumor cells (**section 3.4**). The identified optimal influx/uptake schedules were then applied to a larger sample of *in silico* tissues with initial oxygenation levels close to the experimental data to assess schedules robustness and reproducibility (**section 3.5**).

### 3.1 Generation of *in silico* tissues with a stable oxygen gradient

In general, the patterns in tissue oxygenation, that is, the extent and localization of well-oxygenated vs. hypoxic regions, depend on the number and placement of vessels and both tumor and stromal cells. Therefore, we generated a collection of *in silico* tissues with different vascularity (a fraction of the tissue that is occupied by the vessels), as well as tumor and stromal cellularity (a fraction of the tissue populated by tumor or stromal cells, respectively). In particular, we considered tissue vascularity to be between 0.5% and 5% of the whole tissue area, with increments of 0.5%. The fractions of tissue inhabited by tumor cells was varied between 10% and 95%, and by stromal cells between 5% and 95%, both with increments of 5%. We have ensured that the total of cellular and vascular fractions does not exceed 100% of the tissue area. The locations of all vessels and cells were chosen randomly, and we applied the repulsive force algorithm described above to resolve overlaps between the cells and vessels. In all these simulations, the oxygen influx from each vessel was assumed identical (with *δ*_*V*_(*t*) = 1) and the oxygen uptake by each tumor cell was the same (with *δ*_*T*_(*t*) = 1).

For each *in silico* tissue we applied the diffusion-reaction equation of oxygen kinetics to generate a stable oxygen gradient. As a stability criterion, we compared the L_2_-norm between two consecutive oxygen distributions:

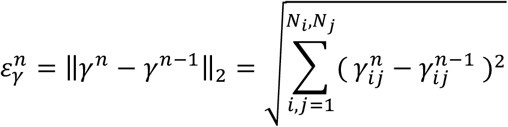

where, 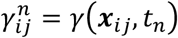 is the oxygen value at the grid point ***x***_*ij*_ at time *t*_*n*_ = *t*_0_ + *n*Δ*t*, and *N*_*i*_ · *N*_*j*_ are the total number of grid points. The oxygen gradient stability was achieved when the normalized error reached a small enough value, i.e., 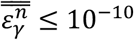, where

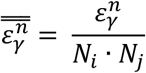

An example of the oxygen gradient stabilization is shown in **Figure 3**. The tissue morphology with 3.5% vascularity, tumor cell fraction of 55%, and stromal cellularity of 30% is presented in **Figure 3A**, and the final oxygen gradient in **Figure 3B**. The initial oxygen level was set up to 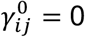 mmHg uniformly in the whole tissue domain and the average oxygen level in the whole tissue stabilized at the level of 29.89 mmHg in about 2×10^4^ iteration steps reaching the normalized error of 9.99×10^−11^ (**Figure 3C**). The final stabilized oxygen level is independent of the oxygen concentration chosen to initiate this process (**Supporting Information S3**).

**Figure 3.**
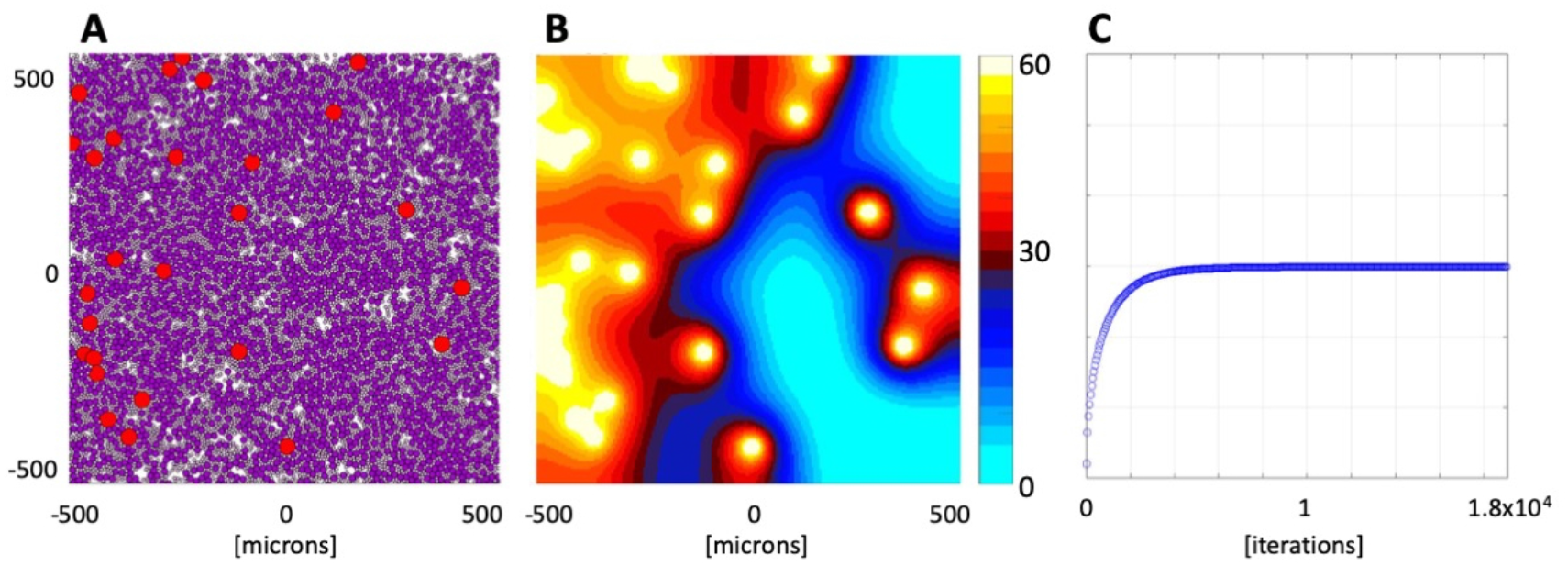
Stabilized oxygen gradient. **A**. *In silico* tissue morphology comprised of 3.5% of vasculature (red circles), 55% of tumor cells (dark purple circles), and 30% of stromal cells (light pink circles). **B**. Tissue oxygenation pattern is color-coded with high oxygen levels (yellow) near the vessels and low oxygen levels (cyan) in poorly vascularized regions. **C**. Changes in the average oxygen concentration over time from initial 0 mmHg to the stable level of 29.89 mmHg.

### 3.2. Classification of tissues with specific saturation levels

The generated library contains 1,530 tumor tissues of different morphologies with a stabilized distribution of oxygen. The minimal and maximal average oxygenations were achieved at the levels of 2.06 mmHg and 55.55 mmHg, respectively. All tumor tissues were divided into five classes according to their average oxygenation (from 0 to 60 mmHg with increments of 12 mmHg).

The parameter space corresponding to each class is shown in **Figure 4A** in a form of a 3D convex hull, that is the smallest convex set containing all data points from a given class (color-coded in blue, red, orange, yellow and white). Almost 1/3 of all tissues (506 cases) have stabilized at the high level of 36-48 mmHg (yellow region). Moderate oxygenation of 24-36 mmHg was achieved in 340 tissues (orange region). Low oxygenation of 12-24 mmHg was reached in 284 tissues (red region), and a similar number of tissues (270) have stabilized on the hypoxic levels of 0-12 mmHg (blue region). The smallest number of tissues (130) reached the very high level of oxygenation above 48 mmHg (white region). In general, the higher tissue vascularity and lower cellularity, the higher level of average tissue oxygenation. However, the vascularity and cellularity values need to be tightly balanced to achieve a desired level of oxygen stability, which is illustrated by three examples in **Figure 4B-D**. Each tissue reached an average oxygenation level near 33 mmHg, however the increased tissue vascularity (from 1.5% to 2.5%, to 4%) is accompanied by increased total tissue cellularity (from 30% to 65%, to 95%). Interestingly, the time required to stabilize the oxygen level is different in each case, with less dense tissues requiring more time to achieve the stable oxygen level (**Figure 4B-D** last column). We also analyzed how the stabilized levels of oxygen depend on the random locations of vessels and cells by generating 25 different tissues with vascularity and cellularity corresponding to those in **Figure 4B-D**. They stabilized at the levels of 32.63, 29.98, and 36.15 mmHg, respectively, with the standard deviation of 2-3 mmHg (**Supporting Information S4**).

**Figure 4.**
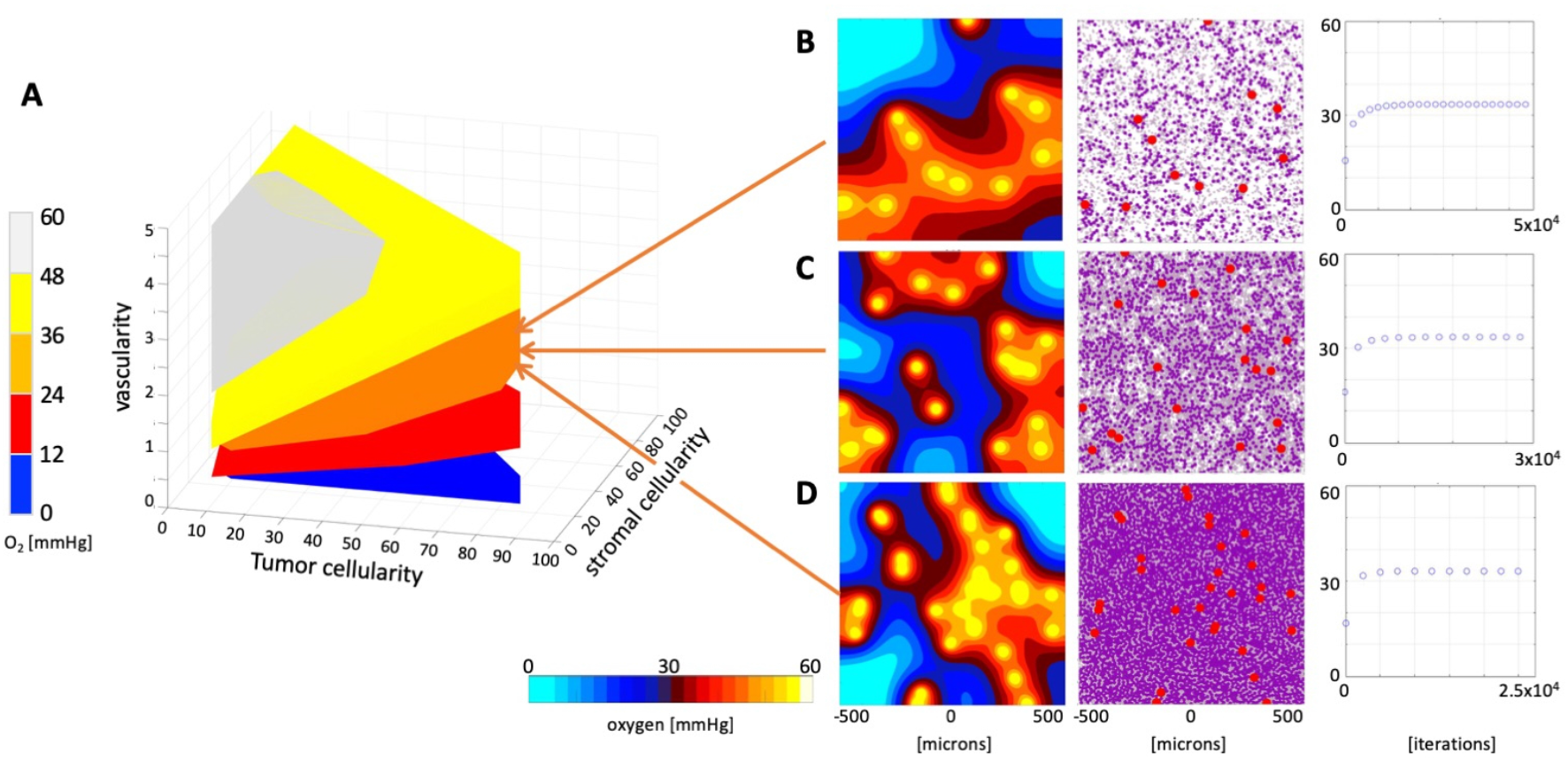
Classification of tissue oxygenation. **A**. A parameter space (convex hulls) of tissues characterized by the vascularity, tumor cellularity and stromal cellularity classified into 5 classes with respect to the stabilized average oxygen level. **B-D**. Three examples of tissues with similar oxygen saturation levels: **B**. A tissue with vascularity 1.5%, tumor cellularity 15%, stromal cellularity 15%, and stable oxygen of 33.43 mmHg. **C**. A tissue with vascularity 2.5%, tumor cellularity 30%, stromal cellularity 35%, and stable oxygen of 33.53 mmHg. **D**. A tissue with vascularity 4%, tumor cellularity 75%, stromal cellularity 20%, and stable oxygen of 33.16 mmHg.

### 3.3. Selection of tissues best-fitted to experimental data

For further analysis we selected computational tissues with stabilized level of oxygen that closely matched the maximal experimental measurement recorded in each of the four ROIs shown in **Figure 1**. In particular, region #1 (black) has initial oxygenation of 36.312 mmHg, and the generated tissue with closest oxygen level of 36.315 mmHg contains vascular fraction of 4%, tumor cell fraction of 15%, and stromal cell fraction of 75% of the tissue area (**Figure 5A**). The experimental region #2 (red) has maximal oxygenation of 22.89 mmHg, and our *in silico* tissue configuration with a vascular fraction of 2.5%, tumor fraction of 45%, and stromal fraction of 40% reached the stable oxygenation of 22.22 mmHg (**Figure 5B**). The third experimental region (blue) has oxygenation of 13.99 mmHg, while the computational tissue with stabilized oxygen of 14.01 mmHg contains vascular fraction of 1.5%, tumor fraction of 40%, and stromal fraction of 45% of the tissue area (**Figure 5C**). Finally, the fourth experimental region (magenta) has maximal oxygenation near 1.83 mmHg, and the closest generated configuration has a stabilized oxygenation of 2.06 mmHg and vascular fraction of 0.5%, tumor fraction of 20%, and stromal fraction of 75% of the tissue area (**Figure 5D**).

**Figure 5.**
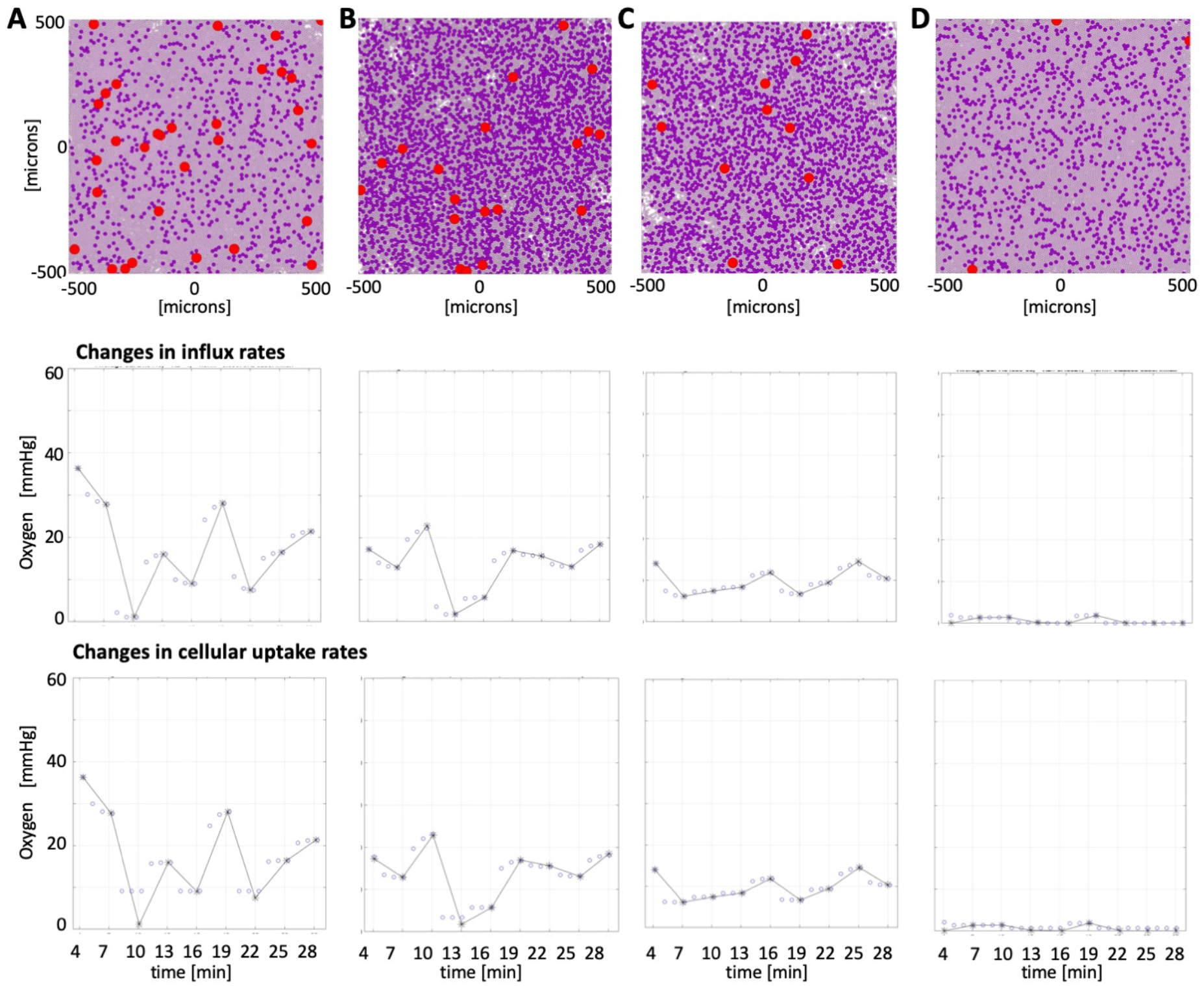
Reconstruction of oxygen fluctuations in ROIs #1-4. Tissue configurations (top row) for which stabilized oxygen levels matched the maximal average oxygenations in each of the region of interests and the reconstructed oxygen fluctuations when either vascular influx rates (middle row) or cellular uptake rates (bottom row) were varied. Open circles denote computational data recorded every 1 minute. Star symbols connected by solid lines denote experimental data recorded every 3 minutes. Tissue characteristics: **A**. ROI#1 (black): vascularity 4%, tumor cellularity 15%, and stromal cellularity 75%. **B**. ROI#2 (red): vascularity 2.5%, tumor cellularity 45%, and stromal cellularity 40%. **C**. ROI#3 (blue): vascularity 1.5%, tumor cellularity 40%, and stromal cellularity 45%. **D**. ROI#4 (magenta): vascularity 0.5%, tumor cellularity 20%, and stromal cellularity 75%.

For each case, our goal was to reproduce oxygen fluctuations shown in **Figure 1C**. Our approach was to alter either the oxygen influx rates or the oxygen cellular uptake rates every three minutes to match each data point that was recorded in *in vivo* experiments. Since the fluctuation in the four selected regions have different magnitudes, this will allow us to assess whether each mechanism may be responsible for the observed changes in tissue oxygen levels. Once these rates were determined for the selected *in silico* tissue, we applied the same schedule to a set of tissues that stabilized at the similar oxygen levels to show how robust these optimal schedules are in reproducing oxygen fluctuations.

### 3.4. Reconstruction of experimentally measured oxygen fluctuation

To test the impact of vascular supply on tissue oxygenation, we simultaneously adjusted the vascular influx of oxygen in each vessel by assigning a fraction *δ*_*V*_(*t*) (between 0 and 1) of the default maximum influx value *γ*_*max*_. To test the role of tumor cell metabolism on tissue oxygenation, we adjusted cellular uptake in all tumor cells by multiplying the default absorption value *T*_*max*_ by a constant ratio *δ*_*T*_(*t*) (between 0 and 50) to account for either decreased or increased cellular uptake. The rates *δ*_*V*_(*t*) and *δ*_*T*_(*t*) were determined using the Mesh Adaptive Direct Search (MADS) method as implemented in MATLAB® by the *patternsearch* routine (22). This optimization algorithm utilizes a value calculated at the given time point by a simulation of the underlying deterministic system and does not require derivatives of the objective function.

Our optimization goal was to minimize the difference between the average tissue oxygenation recorded experimentally, 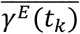, and the one computed by our model, 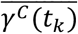, at the end of each 3-minute interval (i.e., for *t*_*k*_ ∈ {4, 7, 10, 13, 16, 19, 22, 25, 28} minutes, *N* = 9 intervals in total), where *x*(*t*_*k-1*_) is a value of either the vascular influx rate *δ*_*V*_(*t*_*k-1*_) or the tumor cell uptake rate *δ*_*T*_(*t*_*k-1*_) at the beginning of each 3-minute time interval (i.e., for *t*_*k-1*_ ∈ {0, 4, 7, 10, 13, 16, 19, 22, 25} minutes):

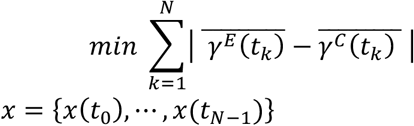

Once the final optimal schedule (influx or uptake) is determined, the normalized L_2_-norm 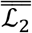 between the simulated and experimental data points is reported as an indication of the goodness of fit (gof) of the optimal schedule:

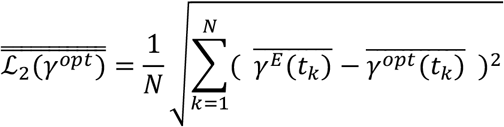

The optimal influx and uptake rates together with the normalized L_2_-norms for all ROIs are listed in the **Table 2**. Method convergence for the cases with best and worse gof (both from ROI#1) is shown in **Supporting Information S5**.

**Table 2.**
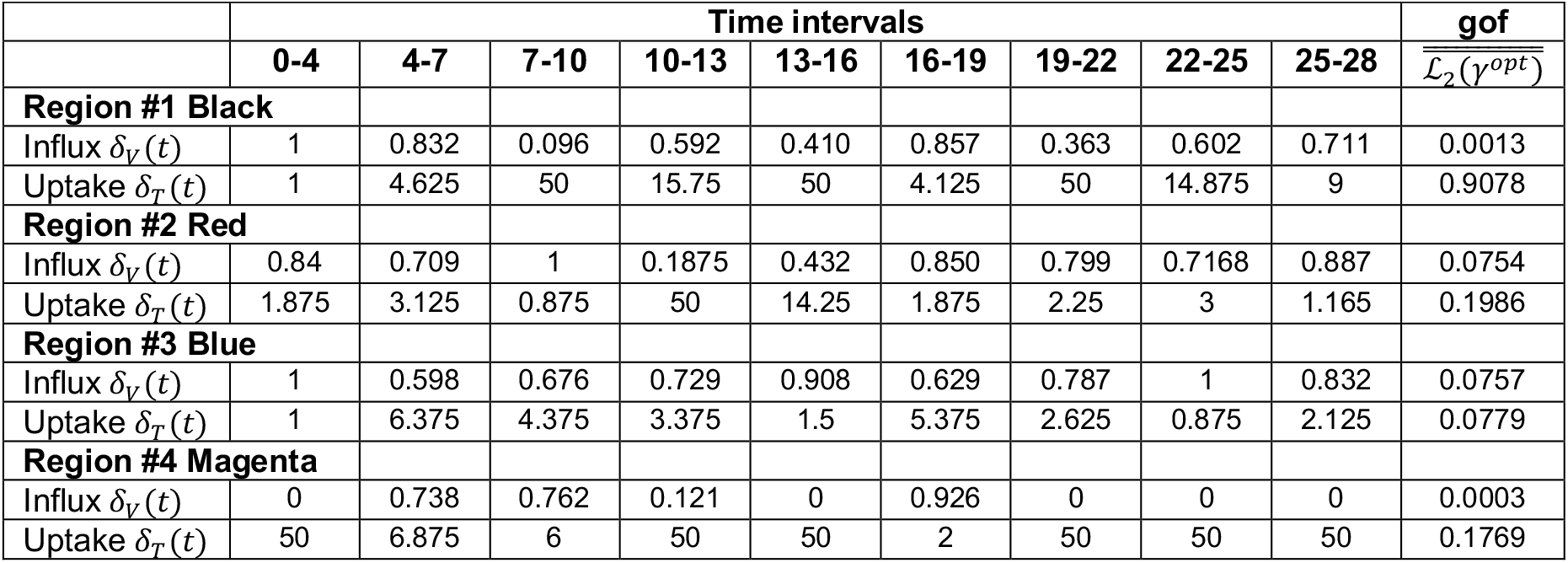
Influx and Uptake Schedules for four considered ROIs.

The resulting simulated fluctuations are shown in **Figure 5**, and the exemplar oxygen distributions at each stage are shown in **Supporting Information S5**. The presented results indicate, that changes in vascular influx can reproduce experimentally observed fluctuations in tissue oxygenation in all four ROIs (in all cases the normalized L_2_-norms are below 0.1). In case of ROI#3, these influx alterations are moderate, not exceeding 40%. However, the sudden drop in tissue oxygenation level observed in ROI#1 and ROI#2 requires a substantial decrease in oxygen influx, as much as 80% and 99%, respectively. In case of ROI#4, when the tissue oxygenation reaches near 0 level, the oxygen influx must be completely shut down. These changes in intravascular oxygenation are physiologically plausible, as the cases with low or even zero arterial oxygen supply (anoxemia) have been observed (23). Here, we showed that short-time fluctuations in tissue oxygenation can be achieved by alterations in intravascular oxygen supply.

The changes in tumor cell metabolisms (modeled as an increase in oxygen uptake) can explain smaller fluctuations in tissue oxygenation (ROI#3, the normalized L_2_-norm below 0.1). It required up to 6-fold changes in the cellular uptake rate to match these fluctuations. This mechanism can also fit cases with near-zero oxygen depletion in the whole tissue patch (ROI#4 and ROI#2, with L_2_-norms near 0.2). However, it failed to reproduce large and rapid fluctuations (ROI#1, with normalized L_2_-norm near 1) even if we considered changes up to 50-fold in cellular uptake. Thus, in general, oxygen fluctuations were not captured by changes in cell metabolism.

### 3.5. Robustness of optimal schedules

The optimal influx/uptake schedules described above were determined using four particular tissues for which the average oxygen level has stabilized the closest to the maximal value recorded for each of the four ROIs in **Figure 1**. Here, we investigated whether these optimal schedules applied to other tissues will reproduce oxygen fluctuations recorded experimentally. One motivation was to test whether tissues of various morphologies but similar average oxygenation levels will respond in a similar way to the schedules that were optimized using one of these tissues. If fluctuations in the average level of tissue oxygenation are not sensitive to tissue morphology, any of these tissues or a small subset of these tissues can be used for further simulation studies of diffusive therapeutic agents and their impact on tumor progression. Second motivation was to provide a link between the average data value recorded for voxels of radiologic images and structure of the tissue that fills the voxel. Potentially, a very large number of tissue structures may result in the same average tissue oxygenation. Our goal here was to investigate whether an additional information, such as temporal data recorded for the same voxel, will result in reduced number of tissue morphologies that reproduce that data. If this tissue number is smaller, we can determine better conditions for selection of tissue morphologies for further studies of intratumoral drug or biomarker distribution. Taken together, we can identify which mechanisms can reproduce the experimentally observed fluctuations, and to provide criteria for selection of different tissues for which these fluctuations can be reproduced.

To achieve our goals, we first identified four sets of tissues with oxygen levels that stabilized within +/- 3.5 mmHg from the maximal value recorded for each ROI (#1-black, #2-red, #3-blue, and #4-magenta). The number of such representative tissues is listed in **Table 3** for each ROI. Next, we applied the optimal influx schedules that were determined for each ROI to the corresponding set of representative tissues. Separately, we also applied the optimal uptake schedules to these tissues. To measure how well the applied schedules reproduced the experimentally observed oxygen fluctuations, we recorded the normalized L_2_-norms 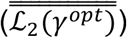, as described above. The total number of considered tissues and the numbers of tissues with L_2_-norms below a threshold value 0.2 are listed in **Table 3** for each ROI.

**Table 3.** Number of tissues representing each ROI and tissues fitting each fluctuation

| Number of representative tissues: | ROI#1 | ROI#2 | ROI#3 | ROI#4 |
| --- | --- | --- | --- | --- |
| within $\pm 3.5$ mmHg from experimental value | 277 | 189 | 147 | 123 |
| with $L_2$-norm$<0.2$ for the influx schedule | 42 | 38 | 40 | 61 |
| with $L_2$-norm$<0.2$ for the uptake schedule | 0 | 6 | 15 | 73 |

The obtained results are also summarized graphically in **Figure 6**. The convex hulls represent a range of tissue characteristics (i.e., vascularity, tumor cellularity, and stromal cellularity) that satisfy a particular condition under consideration. The cyan convex hulls in **Figure 6A** represent all tissues with the oxygen level that stabilized within +/- 3.5 mmHg from the maximal value recorded for each ROI. A subset of tissues for which the optimal influx schedule resulted in oxygen fluctuations that fitted the experimental data with the normalized L_2_-norm below 0.2 are shown as green convex hulls. Similarly, a subset of tissues for which the optimal uptake schedule fitted the experimental data with the L_2_-norm below 0.2 are shown in black. The scatter plots in **Figure 6B** show the relationship between and the initial stable oxygenation of this tissue (x-axis shows the deviation of the tissue initial oxygenation from the experimental measurement for a given ROI) and the normalized L_2_-norm value for each tissue (y-axis). The green dots represent data for optimal influx schedule and grey dots show data for optimal uptake schedule. The red dashed lines represent the normalized L_2_-norm value of 0.2.

**Figure 6.**
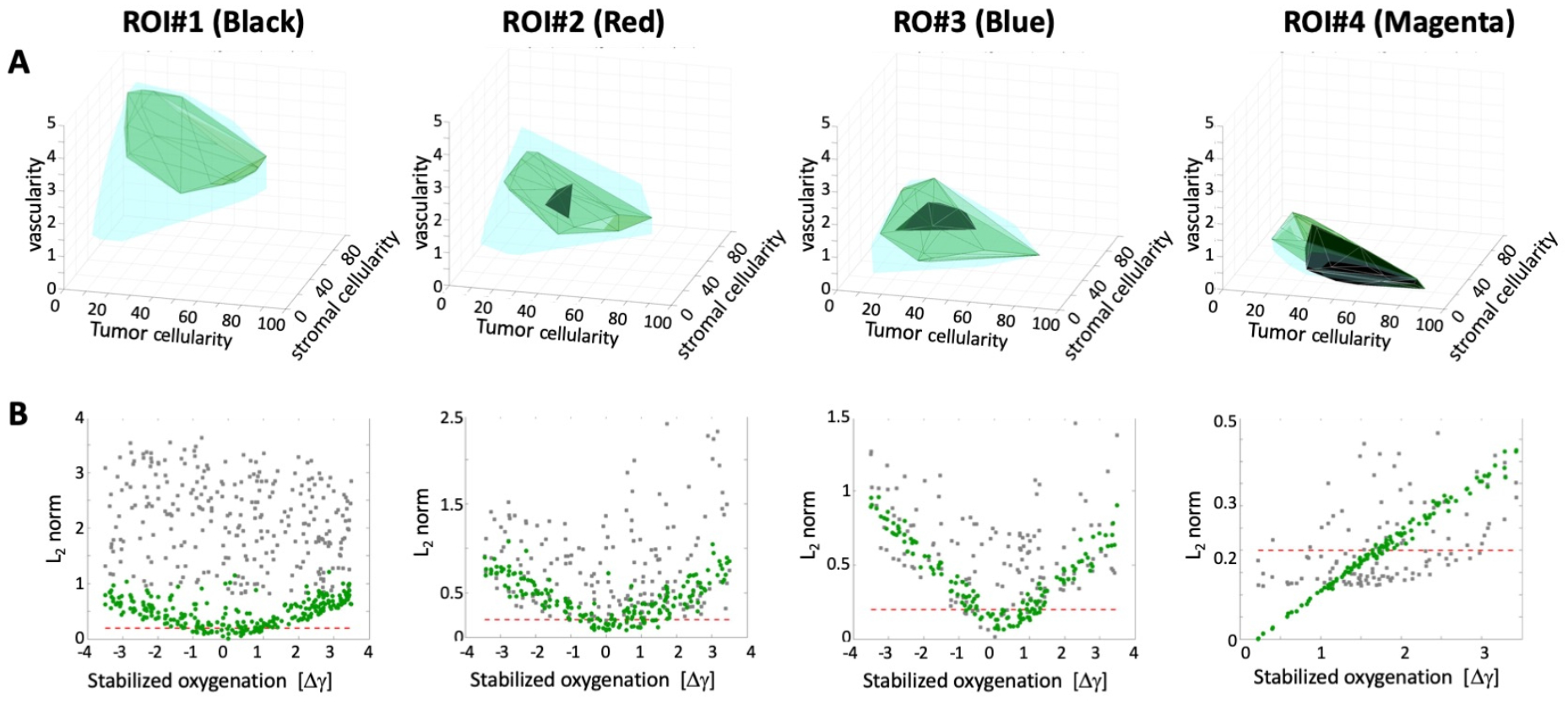
Robustness of optimal influx and uptake schedules. **A**. Parameter spaces of all tissues with oxygen level within +/- 3.5 mmHg from the maximal experimental value for each ROI (3D convex hulls shown in cyan, together with convex hulls for optimal influx schedule (green) and optimal uptake schedule (black) that fit experimental data with normalized L_2_-norm smaller than 0.2. **B**. Normalized L_2_-norms for influx schedule (green dots) and uptake schedule (grey dots) for each tissue from the cyan convex hull. The red dashed line represents the L_2_-norm value of 0.2. The results shown from left to right for: ROI#1 (black), ROI#2 (red), ROI#3 (blue), ROI#4 (magenta).

For each ROI, the optimal influx schedule was most successful for tissues with initial oxygen levels that stabilized near the experimental measurements, since all green dots located below the red threshold line are concentrated near the 0 value in **Figure 6B**. These tissues are also co-localized in **Figure 6A**, though the tissue characteristics shown as the green convex hulls span a broader range of values. The exception is ROI#4, for which almost all considered tissues responded to influx and uptake schedules by following the experimental fluctuations (the green and black convex hulls almost overlap with the cyan one). Though, the fluctuations in this region were very small, thus they were easier to reproduce by both schedules. With the increased magnitude of oxygen fluctuations (from ROI#3 to ROI#2, to ROI#1), the numbers of representative tissues that followed the experimental data was decreased, and there was no tissue in ROI#1 for which the optimal cellular uptake schedule resulted in fitting with the normalized L_2_-norm below 0.2 (no black convex hull).

These results bring several interesting observations. We showed that only a fraction of tissue morphologies with oxygen level that stabilized near the given experimental data are able to reproduce temporal changes in tissue oxygenation when the vascular influx of oxygen is varied. These well-fitted tissues have a stabile oxygen level within +/- 1mmHg from the experimental data. Thus, for future applications we can reduce the searching radius from 3.5 to 1 mmHg in order to find plausible tissue morphologies.

By comparing simulation results with the vascular influx schedule vs. the cellular uptake schedule, we conclude that alterations in vascular levels of oxygen were able to reproduce the observed fluctuations. On the other hand, in order to achieve the same effect when the metabolic changes in tumor cells are considered, the cells would need to increase their oxygen absorption by 50-fold over a period of 3 minutes, which may not be biologically feasible. While it was reported in literature that cellular uptake of oxygen can vary greatly between the cell lines (i.e., 1-350 amol/s per cell (24), and 1-120 amol/s per cell (25)), changes reported in the same cells varied no more than 10-fold when the culture conditions were modified (26, 27).

## 4. Discussion

In this paper, we investigated the mechanisms leading to frequent fast fluctuations in tissue oxygenation observed experimentally in mice models. By using the EPR imaging, it was possible to record differences in average partial oxygen pressure in specific regions of interests (ROIs) within the tumor tissue. These fluctuations were as high as 30 mmHg over a 3 minutes long interval. The first hypothesis we tested was that these fluctuations are related to changes in vascular oxygen supply. However, the same mouse experiments showed no differences in the levels of an imaging tracer in the same ROIs. Therefore, we also tested a hypothesis that modulations in cellular oxygen absorption may be responsible for the observed fluctuations in tissue oxygenation. We first generated a large number of in silico tumor morphologies and selected those for which stable oxygenation levels were the closest to the experimental data in each ROI. Next, we used computational optimization techniques to determine influx and uptake schedules that best fitted the experimental fluctuation data. Finally, we applied these optimal schedules to other tissues with similar oxygenation levels to test whether they will respond in a similar way to the same influx/uptake schedules. This procedure showed that fast changes in vascular supply of oxygen can explain the observed fluctuations in all considered ROIs. However, in order to fit the large magnitude fluctuation data using a modified oxygen uptake schedule, the 50-fold changes in cellular oxygen absorption would have to occur over 3-minute intervals, which may not be biologically feasible. Thus, we showed computationally which mechanisms are more probable to lead to cyclic hypoxia.

Moreover, we provided a link between the average data value recorded for voxels of radiologic images and the cellular/vascular architecture of the tissue that fills these voxels. Since a single voxel of the radiologic image has an area about a millimeter square, the underlying tissue patch is large enough to contain subregions of different characteristics. For example, it may include zones without functional vasculature, with sever hypoxia, or high cellular density poorly penetrated by drugs, or may harbor resistant tumor cell subpopulations. Such small-scale phenomena will not be reflected in the average values reported by radiologic images. To capture information on the cellular scale based on average voxel values novel computational approaches are needed. Potentially, a very large number of tissue architectures may result in the same average oxygenation value. However, by adding a series of temporal data recorded for the same voxel, we showed that the number of tissue morphologies capable of reproducing the underlying temporal dynamics of oxygen in that voxel is greatly reduced. In the cases discussed here, we were left with about 40 different tissues for each ROI out of over 1,500 different initial tissues. This is a computationally manageable number of distinct tissues for any further analysis or simulations, giving us a tool for microscale modeling (on individual cell level) based on macroscale data (average value in tissue voxel).

While we used here the pre-clinical data collected using EPR imaging, our goal for future studies is to develop a similar macro-scale to micro-scale model based on clinically-relevant radiologic imaging. Computed tomography (CT), magnetic resonance imaging (MRI), or positron emission tomography (PET), all provide a way to monitor tumor development and response to therapies in a longitudinal and minimally invasive fashion. Of particular interest is dynamic contrast-enhanced MRI (DCE-MRI), which is a diagnostic imaging method that can capture information on tissue perfusion. In this method, intravenous contrast medium is infused, and images are acquired at multiple time points in order to capture the time-varying signal intensity in each voxel. This temporal intensity information can determine maps of tumor perfusion, microvascular permeability, vascular volume fraction, extracellular-extravascular volume fraction, and diffusivity of tissue water, that combined can predict outcomes and guide therapy (28, 29). To augment temporal resolution, new under-sampling techniques coupled with parallel imaging methods (GRASP, Golden-angle RAdial Sparse Parallel imaging) can be used to continuously acquire images over a long-time continuum (i.e. 5 minutes) at repeatable, small time intervals (i.e. 2.5 seconds), generating a detailed library of exquisite time resolved data. This approach accommodates different attempts to quantify rates of tissue contrast enhancement and to calculate perfusion kinetics (30). Two other non-invasive imaging approaches, TOLD MRI and BOLD MRI, can be used to visualize information on tumor oxygenation and vascular hemodynamics (31). These methods exploit the paramagnetic nature of dissolved oxygen, and the fact that deoxy-hemoglobin is paramagnetic (and therefore MRI active) but oxy-hemoglobin is diamagnetic (and thus MRI-silent). For example, successive sets of BOLD MRI images can be acquired with the subject breathing first standard medical air (21% O_2_), then breathing carbogen (95% O_2_, 5% CO_2_), and finally when breathing hypercapnic air (21% O_2_, 5% CO_2_). The systematic changes in the BOLD MRI signal in each voxel in response to vasodilation and increased blood oxygenation produced by the different inhaled gas mixtures can reveal information about the microvasculature, albeit at a millimeter length scale (32).

The micro-macro scale mechanistic link developed in this paper can be used in the future to generate a set of representative morphologies for cell-scale simulations of various anti-cancer treatments (chemotherapy, hypoxia-activated targeted therapy, radiotherapy, or immunotherapy). These microscopic level predictions of tumor response and specific tracer distribution can then be compared to longitudinal radiologic data on tissue voxel scale. This can give us an insight into heterogeneities of intratumoral drug distributions or immune cell penetration patterns, and the emergence of tissue subregions which can harbor chemo-, radio-, or immune-resistant cells. Thus, the mechanistic models of the tumor, such as we have described here, will enable patient-specific simulations to predict the trajectory of tumor response to specific interventions.

## Supporting Information Captions

**Supporting Information S1. Resolving cell overlapping conditions. Supporting Information S2. Model Parameters**.

**Supporting Information S3. Oxygen stabilization and its dependence on initial oxygen concentration**.

**Supporting Information 4. Analysis of oxygen stabilization for tissues of identical characteristics but different morphologies**.

**Supportive Information S5. MABS method convergence for influx and uptake schedules**.

## Author Contributions

This project was conceptualized by KAR, JLK, NR, and JRC. Computer simulations, visualization and formal analysis were completed by JLK. The methodology was developed by JLK and KAR. The manuscript was written by JLK, KAR, NR and JRC.

## Acknowledgment

This work was supported in part by the U01-CA202229 Physical Sciences Oncology Project (PS-OP) grant from the US National Institutes of Health, National Cancer Institute (to KR), the Moffitt Radiology Pilot Project grant (to JC), and Shared Resources at the H. Lee Moffitt Cancer Center & Research Institute an NCI designated Comprehensive Cancer Center (Moffitt IRAT Core) under the grant P30-CA076292 from the National Institutes of Health).

## References

1. Harris LH. Hypoxia-a key to rgulatory factor in tumour growth. Nature Reviews Bacer. 2002;2:38–47.

2. Saxena K, Jolly K. Acute vs. Chrinic vs. Cyclic Hypoxia: Their differentual dynamics, molecular mechanisnsm and effects on tumor progression. Biomolecules. 2019;9:339.

3. Dewhirst M, Cao Y, Moeller B. Cycling hypoxia and free radicals regulate angiogenesis and radiotherapy response Nat Rev Cnacer. 2008;8(6):425–37.

4. Michiels C, Tellier C, Feron O. Cuncling hypoxia: a key feature of the tumor microenvironment. Biochimica et Biophysica Acta. 2016;1866:76–86.

5. Yasui H, Matsumoto S, Devasahayam N, Munasinghe JP, Choudhuri R, Saito K, Subramanian S, Mitchell JB, Krishna MC. Low-field magnetic resonance imaging to visualize chronic and cycling hypoxia in tumor-bearing mice. Cancer Res. 2010;70(16):6427–36. doi: 10.1158/0008-5472.CAN-10-1350. PubMed PMID: 20647318; PMCID: PMC2922437.

6. Matsuo M, Matsumoto S, Mitchell JB, Krishna MC, Camphausen K. Magnetic resonance imaging of the tumor microenvironment in radiotherapy: perfusion, hypoxia, and metabolism. Semin Radiat Oncol. 2014;24(3):210–7. Epub 2014/06/17. doi: 10.1016/j.semradonc.2014.02.002. PubMed PMID: 24931096; PMCID: PMC4060050.

7. Elas M, Bell R, Hleihel D, Barth ED, McFaul C, Haney CR, Bielanska J, Pustelny K, Ahn KH, Pelizzari CA, Kocherginsky M, Halpern HJ. Electron paramagnetic resonance oxygen image hypoxic fraction plus radiation dose strongly correlates with tumor cure in FSa fibrosarcomas. Int J Radiat Oncol Biol Phys. 2008;71(2):542–9. Epub 2008/05/14. doi: 10.1016/j.ijrobp.2008.02.022. PubMed PMID: 18474313; PMCID: PMC2577780.

8. Christodoulou AG, Redler G, Clifford B, Liang ZP, Halpern HJ, Epel B. Fast dynamic electron paramagnetic resonance (EPR) oxygen imaging using low-rank tensors. J Magn Reson. 2016;270:176–82. Epub 2016/08/09. doi: 10.1016/j.jmr.2016.07.006. PubMed PMID: 27498337; PMCID: PMC5127203.

9. Karolak A, Poonja S, Rejniak KA. Morphophenotypic classification of tumor organoids as an indicator of drug exposure and penetration potential. PLoS Comput Biol. 2019;15(7):e1007214. Epub 2019/07/17. doi: 10.1371/journal.pcbi.1007214. PubMed PMID: 31310602; PMCID: PMC6660094.

10. Perez-Velazquez J, Gevertz JL, Karolak A, Rejniak KA. Microenvironmental Niches and Sanctuaries: A Route to Acquired Resistance. Adv Exp Med Biol. 2016;936:149–64. Epub 2016/10/16. doi: 10.1007/978-3-319-42023-3_8. PubMed PMID: 27739047; PMCID: PMC5113820.

11. NCI-60 Human Tumor Cell Lines Screen [Internet]2015. Available from: https://dtp.cancer.gov/discovery_development/nci-60/.

12. Shashni B, Ariyasu S, Takeda R, Suzuki T, Shiina S, Akimoto K, Maeda T, Aikawa N, Abe R, Osaki T, Itoh N, Aoki S. Size-Based Differentiation of Cancer and Normal Cells by a Particle Size Analyzer Assisted by a Cell-Recognition PC Software. Biol Pharm Bull. 2018;41(4):487–503. doi: 10.1248/bpb.b17-00776. PubMed PMID: 29332929.

13. Caramalho I, Faroudi M, Padovan E, Müller S, Valitutti S. Visualizing CTL/melanoma cell interactions: multiple hits must be delivered for tumour cell annihilation. J Cell Mol Med. 2009;13(9B):3834–46.

14. Rieger H, Fredrich T, Welter M. Physics of the tumor vasculature: Theory and experiment. Eur Phys J Plus. 2016;131:31.

15. Baumgartner W, Drenckhahn D. Transmembrane cooperative linkage in cellular adhesion. Eur J Cell Biol. 2002;81(3):161–8. doi: 10.1078/0171-9335-00233. PubMed PMID: 11998868.

16. Rejniak K, Dillon RH. A single cell-based model of the ductal tumour microarchitecture. Computational and Mathematical Methods in Medicine. 2007;8(1):51–69.

17. Shimolina LE, Izquierdo MA, Lopez-Duarte I, Bull JA, Shirmanova MV, Klapshina LG, Zagaynova EV, Kuimova MK. Imaging tumor microscopic viscosity in vivo using molecular rotors. Sci Rep. 2017;7:41097. doi: 10.1038/srep41097. PubMed PMID: 28134273; PMCID: PMC5278387.

18. Latour LL, Svoboda K, Mitra PP, Sotak CH. Time-dependent diffusion of water in a biological model system. Proc Natl Acad Sci U S A. 1994;91(4):1229–33. doi: 10.1073/pnas.91.4.1229. PubMed PMID: 8108392; PMCID: PMC43130.

19. Schornack PA, Gillies RJ. Contributions of cell metabolism and H+ diffusion to the acidic pH of tumors. Neoplasia. 2003;5(2):135–45. doi: 10.1016/s1476-5586(03)80005-2. PubMed PMID: 12659686; PMCID: PMC1502399.

20. Jain RK. Determinants of tumor blood flow: a review. Cancer Res. 1988;48(10):2641–58. Epub 1988/05/15. PubMed PMID: 3282647.

21. Casciari J, Sotirchos S, Sutherland R. Variations in tumor cell growth rates and metabolism with oxygen concentration, glucose concentration, and extracellular pH.1992;151(2):386–94.

22. Venkataraman P. Applied Optimization with MATLAB Programming Wiley; 2009.

23. Sarkar M, Niranjan N, Banyal PK. Mechanisms of hypoxemia. Lung India. 2017;34(1):47–60. Epub 2017/02/02. doi: 10.4103/0970-2113.197116. PubMed PMID: 28144061; PMCID: PMC5234199.

24. Wagner BA, Venkataraman S, Buettner GR. The rate of oxygen utilization by cells. Free Radic Biol Med. 2011;51(3):700–12. Epub 2011/06/15. doi: 10.1016/j.freeradbiomed.2011.05.024. PubMed PMID: 21664270; PMCID: PMC3147247.

25. Place TL, Domann FE, Case AJ. Limitations of oxygen delivery to cells in culture: An underappreciated problem in basic and translational research. Free Radic Biol Med. 2017;113:311–22. Epub 2017/10/17. doi: 10.1016/j.freeradbiomed.2017.10.003. PubMed PMID: 29032224; PMCID: PMC5699948.

26. Wojtkowiak JW, Cornnell HC, Matsumoto S, Saito K, Takakusagi Y, Dutta P, Kim M, Zhang X, Leos R, Bailey KM, Martinez G, Lloyd MC, Weber C, Mitchell JB, Lynch RM, Baker AF, Gatenby RA, Rejniak KA, Hart C, Krishna MC, Gillies RJ. Pyruvate sensitizes pancreatic tumors to hypoxia-activated prodrug TH-302. Cancer Metab. 2015;3(1):2. Epub 2015/01/31. doi: 10.1186/s40170-014-0026-z. PubMed PMID: 25635223; PMCID: PMC4310189.

27. Rose S, Frye RE, Slattery J, Wynne R, Tippett M, Pavliv O, Melnyk S, James SJ. Oxidative stress induces mitochondrial dysfunction in a subset of autism lymphoblastoid cell lines in a well-matched case control cohort. PLoS One. 2014;9(1):e85436. Epub 2014/01/15. doi: 10.1371/journal.pone.0085436. PubMed PMID: 24416410; PMCID: PMC3885720.

28. Stringfield O, Arrington JA, Johnston SK, Rognin NG, Peeri NC, Balagurunathan Y, Jackson PR, Clark-Swanson KR, Swanson KR, Egan KM, Gatenby RA, Raghunand N. Multiparameter MRI Predictors of Long-Term Survival in Glioblastoma Multiforme. Tomography. 2019;5(1):135–44. Epub 2019/03/12. doi: 10.18383/j.tom.2018.00052. PubMed PMID: 30854451; PMCID: PMC6403044.

29. Lorza AMA, Ravi H, Philip RC, Galons JP, Trouard TP, Parra NA, Von Hoff DD, Read WL, Tibes R, Korn RL, Raghunand N. Dose-response assessment by quantitative MRI in a phase 1 clinical study of the anti-cancer vascular disrupting agent crolibulin. Sci Rep. 2020;10(1):14449. Epub 2020/09/04. doi: 10.1038/s41598-020-71246-w. PubMed PMID: 32879326; PMCID: PMC7468301.

30. Feng L, Wen Q, Huang C, Tong A, Liu F, Chandarana H. GRASP-Pro: imProving GRASP DCE-MRI through self-calibrating subspace-modeling and contrast phase automation. Magn Reson Med. 2020;83:94–108.

31. Hallac RR, Zhou H, Pidikiti R, Song K, Stojadinovic S, Zhao D, Solberg T, Peschke P, Mason RP. Correlations of noninvasive BOLD and TOLD MRI with pO2 and relevance to tumor radiation response. Magnetic resonance in medicine. 2014;71(5):1863–73. Epub 2013/07/03. doi: 10.1002/mrm.24846. PubMed PMID: 23813468; PMCID: PMC3883977.

32. Landowski TH, Guntle GP, Zhao D, Jagadish B, Mash EA, Dorr RT, Raghunand N. Magnetic Resonance Imaging Identifies Differential Response to Pro-Oxidant Chemotherapy in a Xenograft Model. Transl Oncol. 2016;9(3):228–35. Epub 2016/05/17. doi: 10.1016/j.tranon.2016.04.007. PubMed PMID: 27267841.

