## Supplemental Material for "Bridging cell-scale simulations and radiologic images to explain short-time intratumoral oxygen fluctuations"

### Supporting Information

#### Supporting Information S1. Resolving cell overlapping conditions.

Tissue morphology is generated by randomly placed tumor cells, stromal cells and vessels within the tissue domain. To resolve the accidental overlapping between the cells, they are subjected to repulsive forces until their positions reach equilibrium. A sequence of images showing how the cell overlapping conditions were resolved is presented in Supporting Figure S1.

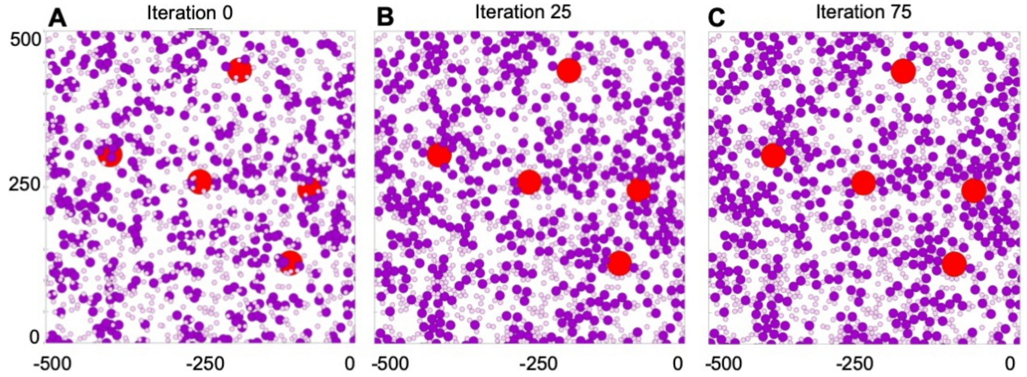


**Supporting Figure S1. Resolving cell overlapping conditions using repulsive forces.** A left-top quarter of the tumor tissue domain characterized by 2% vascularity, 30% tumor cellularity, and 15% stromal cellularity with tumor cells represented by purple circles, stromal cells as pink circles, and vessels as red circles. **A.** An initial iteration 0 before repulsive forces are applied, showing overlapping cells and vessels. **B.** By iteration 25, repulsive forces have been applied and cell relocation has begun; only some cells remained overlapped. **C.** In iteration 75, all overlapping conditions have been resolved and cell positions have stabilized to reflect no overlapping.

#### Supporting Information S2. Model Parameters.

This model represents features characteristic of many solid tumors, thus we did not focus on any specific tumor type. All parameter ranges used in our simulations are based on scientific literature. Solid tumor cells have typically diameter between 10 and 25 microns (1) and we used the middle value of $2R_T = 15\,\mu m$ for tumor cells in the model. Tumor stoma contains various non-tumor cells, such as fibroblasts, lymphocytes, microphages, or dendritic cells. These cells are smaller than tumor cells, and typically have sizes between 6 and 10 microns (2). We used the intermediate value of $2R_S = 7.5\ \mu m$ for all stromal cells in our simulations. Tumor tissue can harbor vasculature of various sizes and oxygen content, from capillaries of 4 microns in diameter and with 30 mmHg of partial oxygen pressure ($pO_2$), to arteries of 50 microns in diameter and $pO_2$ of 80 mmHg (3). We used the vascular size of $2R_V = 40\ \mu m$ for all blood vessels in our simulations and the base oxygen content in each vessel was set up to 60 mmHg. However, to unify all units in the model, we used the scaling factor of $\sigma = 0.5 \times 10^{-19}$ giving the base vascular oxygen level of $\gamma_{max} = 60\ \sigma g/\mu m^3$. In simulations, when the optimal oxygen influx schedules were determined, we allowed the vascular oxygen level to vary between $0\ \sigma g/\mu m^3$ and $60\ \sigma g/\mu m^3$, $(0 \leq \delta_V(t) \leq 1)$ simultaneously in each vessel. The mean experimental value of oxygen diffusion in pure water at 20°C is of order of $10^{-5} cm^2/s$, however the effective diffusion coefficient in tumor tissue is slower, of the order of $\mathcal{D}_\gamma = 10^{-6} cm^2/s = 100 \mu m^2/s$ (4). Oxygen uptake by both tumor and stromal cells is modeled using the Michaelis-Menten equation with parameters values estimated in (5). We used a common Michaelis constant: $\kappa_m = 2.1 \times 10^{-3}\ mM$ of $O_2$ (=$1.134\ \sigma g/\mu m^3$ after scaling), and the maximum uptake rate of: $6.25 \times 10^{-17}\ mole/s$ per cell (=$0.382\ \sigma g/s \cdot \mu m^3$ per grid, after scaling). Thus, the $S_{max} = 0.382\ \sigma g/(s \cdot \mu m^3) \times \pi R_S{}^2$, and $T_{max} = 0.382\ \sigma g/(s \cdot \mu m^3) \times \pi R_T{}^2$. In simulations, when the optimal oxygen uptake schedules were determined, we allowed the cellular uptake level to vary between $0\ \sigma g/(s \cdot \mu m^3)$ and $19.1\ \sigma g/(s \cdot \mu m^3) \times \pi R_T{}^2$, $(0 \leq \delta_V(t) \leq 50)$ simultaneously in each tumor cell. The experimentally measured viscosity of extracellular collagen in tumor tissues is equal to $260 \pm 15\ cP$, and we used this average value as the viscosity of tumor interstitium that we model as a homogeneous gel ($\nu = 250\ \mu g/\mu m \cdot s$ after scaling). The stiffness of adhesive bonds between the cells was reporter to be in a range of $0.1 - 1000 g/cm \cdot s^2$ (6, 7), so in order to separate the overlapping cells and vessels, we used $\mathcal{F} = 50\ \mu g/\mu m \cdot s^2$ (after scaling) as a stiffness of all repulsive forces.

#### Supporting Information S3. Oxygen stabilization and its dependence on initial oxygen concentration.

Stabilization of oxygen distribution within the tissue is a balance between constant oxygen supply from all vessels and constant oxygen uptake by all tumor and stromal cells located within that tissue. Each tissue is generated with a set level of oxygen distributed uniformly throughout the domain. Over time, the oxygen gradient will stabilize within the tissue, as vessels will continue outputting oxygen at a constant rate, and all cells will uptake oxygen according to Michealis-Menten equation. The oxygen gradient stability is achieved when it does not change significantly between consecutive iterations, i.e., when the difference between two consecutive oxygen distributions reaches a small threshold value. Several snapshots showing stabilization of oxygen gradient within the tissue are shown in Supporting Figure S3. The obtained stable oxygen gradient is independent of the initial oxygen concentration. It has been confirmed by running the stabilization routine with the uniform initial oxygen concentration between 0% and 100% of the maximum possible oxygen concentration of 60 mmHg, with the increments of 5%. The results are summarized in Supporting Table 1, and in Supporting Figure S3.

| Initial $O_2$ | 0% | 5% | 10% | 15% | 20% | 25% |
|---|---|---|---|---|---|---|
| Stabilized $O_2$ | 29.894673 | 29.894673 | 29.894673 | 29.894673 | 29.894673 | 29.894674 |
| **Initial $O_2$** | **30%** | **35%** | **40%** | **45%** | **50%** | **55%** |
| **Stabilized $O_2$** | 29.894677 | 29.894694 | 29.894716 | 29.894719 | 29.894720 | 29.894720 |
| **Initial $O_2$** | **60%** | **65%** | **70%** | **75%** | **80%** | **85%** |
| **Stabilized $O_2$** | 29.894720 | 29.894720 | 29.894720 | 29.894720 | 29.894720 | 29.894720 |
| **Initial $O_2$** | **90%** | **95%** | **100%** | **average +/- std** | | |
| **Stabilized $O_2$** | 29.894720 | 29.894720 | 29.894720 | 29.894703 +/- 2.202e-05 | | |

**Supporting Table 1: Initial tissue oxygenation vs. final stabilized oxygen level.** Stabilized average oxygen levels for tissue of characteristics: 3.5% vascular fraction, 55% tumor cellularity, and 30% stromal cellularity, and initial uniform oxygen concentration reported as % of the maximal value of 60 mmHg. The stabilized oxygen levels units: mmHg.

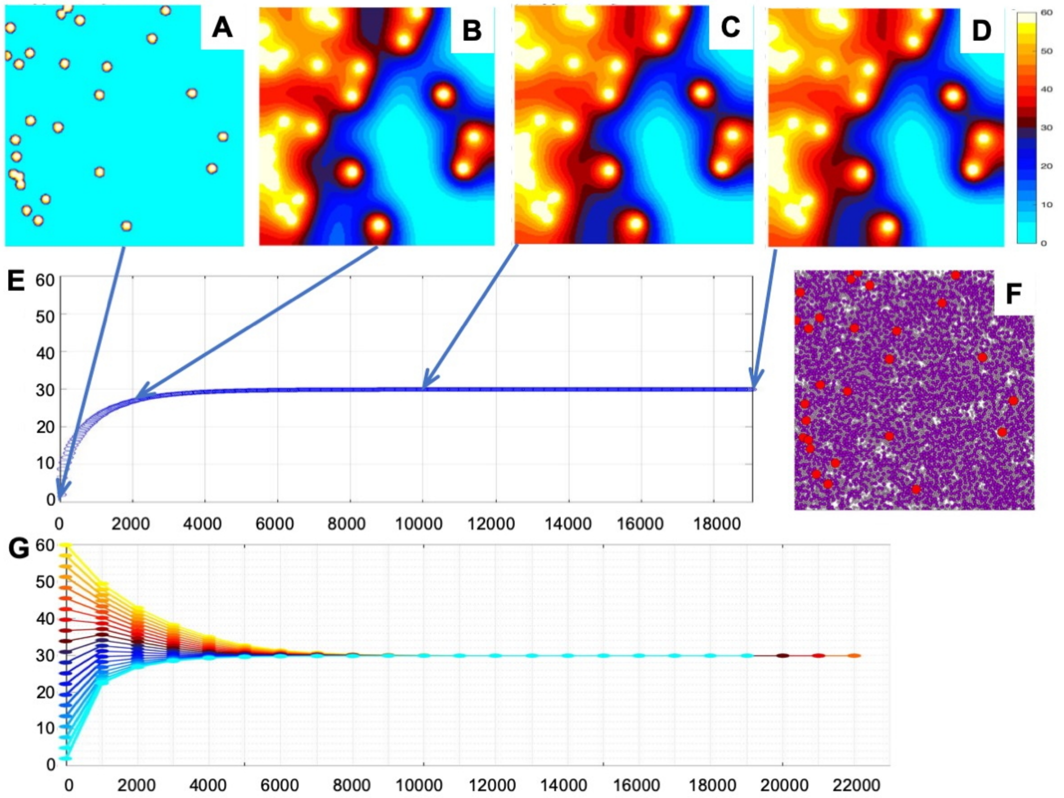


**Supporting Figure S3. Oxygen stabilization within the tumor tissue and the role of initial oxygen concentration. A-D.** Snapshots showing oxygen distribution during the stabilization process, at iterations 0, 2000, 10000, and 18970. **E.** Temporal evolution of the average oxygen level from 0 mmHg until is stabilizes at the 29.89 mmHg with the stabilization error below $10^{-10}$. **F.** Tissue morphology with 3.5% vascular fraction, 55% tumor cellularity, and 30% stromal cellularity; vessels are represented by red circles, tumor cells by purple circles, and stromal cells by pink circles. **G.** Temporal evolution of average oxygen levels for 21 simulations of the same tissue shown in **F.** Each simulation is indicated by a different color that corresponds to initial uniform tissue oxygenation. All simulations stabilized around the 29.89 mmHg, however, simulations with lower initial oxygen concentrations have stabilized faster (cyan and blue lines) than those that started with higher initial oxygen concentrations (red and yellow lines).

#### Supporting Information 4. Analysis of oxygen stabilization for tissues of identical characteristics but different morphologies.

To analyze whether the stabilized levels of oxygen depend on the random locations of vessels and cells, we generated 25 tissues of the same characteristics corresponding to those in Figure 4B-D but different locations of vessels and cells. In particular, we considered three sets of tissues: (a) with 1.5% vascularity, 15% tumor cellularity, and 15% stromal cellularity (Supporting Figure S4.A,D-E) that stabilized at the 32.63 mmHg in average, with standard deviation of 3.3 mmHg; (b) with 2.5% vascularity, 30% tumor cellularity, and 35% stromal cellularity (Supporting Figure S4.B,F-G) that stabilized at the 29.98 mmHg in average, with standard deviation of 2.4 mmHg; and (c) with 4% vascularity, 75% tumor cellularity, and 20% stromal cellularity (Supporting Figure S4.C,H-I) that stabilized at the 36.15 mmHg in average, with standard deviation of 2.4 mmHg. The stabilized oxygen levels for each tissue together with oxygen distribution in the cases of minimal and maximal stabilized oxygen levels are shown in Supporting Figure S4.

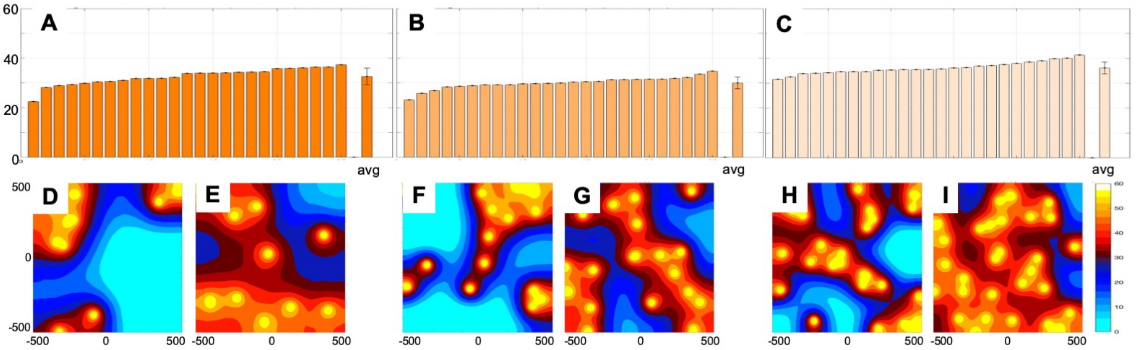


**Supporting Figure S4. Analysis of oxygen stabilization for tissues of identical characteristics but different morphologies. A**. Levels of oxygen stabilization for 25 tissues of vascularity: 1.5%, tumor cellularity: 15%, stromal cellularity: 15%, with an average oxygenation level of 32.63 mmHg +/- 3.3 mmHg. **B**. Levels of oxygen stabilization for 25 tissues of vascularity: 2.5%, tumor cellularity: 30%, stromal cellularity: 35%, with an average oxygenation level of 29.98 mmHg +/- 2.4 mmHg. **C**. Levels of oxygen stabilization for 25 tissues of vascularity: 4%, tumor cellularity: 75%, stromal cellularity: 20%, with an average oxygenation level of 36.15 mmHg +/- 2.4 mmHg. **D**. Spatial gradient of oxygen with lowest average level among tissues in **A. E-I**. Spatial gradients of oxygen with highest (left) and lowest (right) average levels among tissues considered in **A-C.**

#### Supportive Information S5. MABS method convergence for specific influx and uptake schedules

The optimal influx and uptake schedules were determined using the Mesh Adaptive Direct Search (MADS) method as implemented in MATLAB® by the *patternsearch* routine. The schedule consists of nine segments, each 3 minutes long. The optimization routine for each segment starts with the oxygen distribution that fits the average experimental data at the beginning of the time segment. Next, the influx/uptake rates are selected and apply in the computational model that simulates tissue oxygenation for a period corresponding to 3 minutes. The final average oxygenation level is then compared to experimental data at the end of this time segment, and new influx/uptake rates are selected. These steps are repeated until the convergence criteria are met; these include: (i) the difference between the current value of the objective function and the value at the previous best influx/uptake

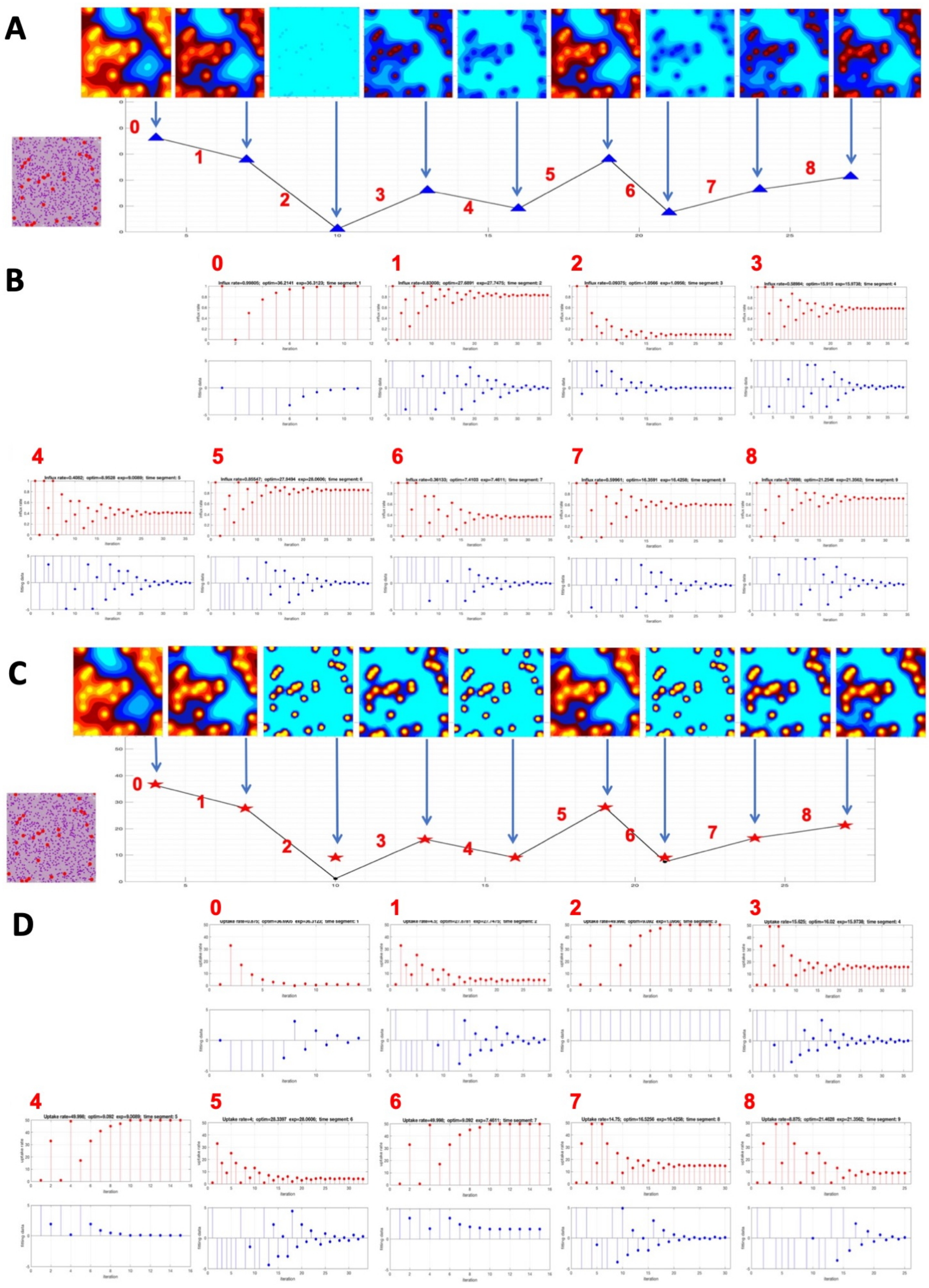


**Supporting Figure S5. Convergence of the MABS method for influx and uptake schedules for ROI #1 (black). A**. Experimental fluctuations (solid line) and simulated fluctuations (blue triangles) for the influx schedule with corresponding oxygen distributions at each time point (above the arrows). An inset shows tissue morphology. **B**. The convergence graphs for each of the 8 time segments with the converging influx rates (red pins) and the converging objective function values (blue pins). **C**. Experimental fluctuations (solid line) and simulated fluctuations (red stars) for the uptake schedule with corresponding oxygen distributions at each time point (above arrows). **D**. The convergence graphs for each of the 8 time segments with the converging uptake rates (red pins) and the converging objective function values (blue pins).

rate is below the prescribed threshold (*FunctionTolerance* of $10^{-3}$), (ii) the difference between influx/uptake rate values at two consecutive iterations is below the prescribed threshold (*StepTolerance* of $10^{-3}$), and (iii) the mesh size determined by the *patternsearch* algorithm is below the threshold (*MeshTolerance* of $10^{-3}$). The objective function is defined as the difference between the simulated average oxygenation and the experimental data. Supporting Figure S5 shows two optimized schedules for influx (Supporting Figure S5AB) and uptake (Supporting Figure S5CD) for the ROI #1 (black). These are the schedules with the best $(\overline{\overline{\mathcal{L}_2(\gamma^{opt})}} = 0.0013)$ and the $(\overline{\overline{\mathcal{L}_2(\gamma^{opt})}} = 0.978)$ goodness of fit, respectively. Supporting Figure S5A shows the experimental data at nine time points (connected by a solid line), the simulated data at the same data points (blue triangles), and oxygen distribution at the same time points (above the arrows). The tissue morphology for which the initial tissue oxygenation was the closest to the experimental data is shown in the inset. The tissue includes vasculature (red circles), tumor cells (purple circles), and stromal cells (pink circles). MABS algorithm convergence at each of the eight time segments is shown in Supportive Figure S5B with influx rate shown by red pins, and the value of the objective function (the difference between the simulated average tissue oxygenation and the experimental one) by blue pins. Note the convergence of both of these values in all panels. The number of MABS algorithm iterations is different for each panel. Supporting Figure S5C shows the experimental data at nine time points (connected by a solid line), the simulated data at the same data points (red stars), and oxygen distribution at the same time points (above the arrows). The tissue morphology is shown in the inset. MABS algorithm convergence at each of the eight time segments is shown in Supportive Figure S5D with uptake rate shown by red pins, and the value of the objective function by blue pins. Note the lack of convergence in panel 2 which corresponds to the simulated data at the end of segment 2 in Supportive Figure S5C being well above the experimental data. This contributes to the high value of the L2-norm.

#### References


1. NCI-60 Human Tumor Cell Lines Screen [Internet]2015. Available from: https://dtp.cancer.gov/discovery_development/nci-60/.
2. Caramalho I, Faroudi M, Padovan E, Müller S, Valitutti S. Visualizing CTL/melanoma cell interactions: multiple hits must be delivered for tumour cell annihilation. J Cell Mol Med. 2009;13(9B):3834-46.
3. Rieger H, Fredrich T, Welter M. Physics of the tumor vasculature: Theory and experiment. Eur Phys J Plus. 2016;131:31.
4. Latour LL, Svoboda K, Mitra PP, Sotak CH. Time-dependent diffusion of water in a biological model system. Proc Natl Acad Sci U S A. 1994;91(4):1229-33. doi: 10.1073/pnas.91.4.1229. PubMed PMID: 8108392; PMCID: PMC43130.

5. Casciari J, Sotirchos S, Sutherland R. Variations in tumor cell growth rates and metabolism with oxygen concentration, glucose concentration, and extracellular pH.1992;151(2):386-94.
6. Baumgartner W, Drenckhahn D. Transmembrane cooperative linkage in cellular adhesion. Eur J Cell Biol. 2002;81(3):161-8. doi: 10.1078/0171-9335-00233. PubMed PMID: 11998868.
7. Rejniak K, Dillon RH. A single cell-based model of the ductal tumour microarchitecture. Computational and Mathematical Methods in Medicine. 2007;8(1):51-69.
